# Environmental drivers define contrasting microbial habitats, diversity, and community structure in Lake Baikal, Siberia

**DOI:** 10.1101/605899

**Authors:** Paul Wilburn, Kirill Shchapov, Edward C. Theriot, Elena Litchman

## Abstract

Understanding how microbial communities respond to environmental change requires knowing the main drivers of their structure, diversity and potential resilience. Lake Baikal is the world’s most ancient, deep, voluminous, and biodiverse lake, holding 20 percent of unfrozen fresh water that is undergoing rapid warming. Despite its global importance, little is known about Baikal’s bacterioplankton communities and their drivers. In this extensive survey, we show that temperature, along with stratification, nutrients, and dissolved oxygen, but not geographic distance, define major microbial habitats and community similarity. Mixed layer and deep water communities exhibited contrasting patterns of richness, diversity and evenness, and comprised different cohesive modules in the whole Baikal OTU co-occurrence network. The network’s small-world properties indicated likely resistance to perturbations but sensitivity to abundance changes of central, most connected OTUs. Previous studies showed Baikal water temperature rising by over 1.2°C since 1946, and this trend is predicted to accelerate. Because temperature emerged as the most significant predictor of the mixed layer community structure, we hypothesize that it is most likely to drive future community changes. Understanding how temperature and other abiotic factors structure microbial communities in this and other rapidly changing ecosystems will allow better predictions of ecosystem responses to anthropogenic stressors.

## INTRODUCTION

The ecological importance of microorganisms in aquatic systems has been recognized at least since the appearance of “ooze” in Lindeman’s trophic energy transfer diagram [1]. Their central place in material and energy fluxes is now recognized for nearly all nutrient cycles, with greater relative importance of prokaryotic organisms in more oligotrophic systems [2]. The advent of next-generation sequencing, combined with environmental monitoring, enabled new discoveries of microbial diversity and function in various aquatic habitats. However, the environmental drivers of microbial community diversity, community structure, function, and stability remain poorly characterized in many aquatic ecosystems, including the world’s most ancient (25 My) lake Baikal – a UNESCO heritage site and known hotspot for endemism of its biota. Baikal is the world’s deepest (1643 m) and most voluminous lake, holding about 20% of world’s surface unfrozen freshwater [3]. Of the approximately 2600 plant and animal species in the lake, two-thirds are endemic, including the dominant primary producers, phytoplankton, zooplankton grazers, benthic and pelagic fish and the top predator – world’s only freshwater seal [3, 4].

Hampton *et al.* [4] showed that Baikal water temperatures have risen by 1.2 °C over 60 years in a high-resolution time series, contributing to an increase in the abundance of non-endemic zooplankton and algal species, with potential consequences for nutrient cycling, food web structure [3] and microbial communities. Moreover, Lake Baikal region is predicted to warm by 3-4°C in the next century [5], with ongoing changes likely to continue and even accelerate. Because the biota of Lake Baikal, including microbial communities, are adapted to cold temperatures, it may be especially vulnerable to warming. Additionally, other changing environmental factors may also alter the lake’s microbial communities.

Despite the remarkable size of this freshwater ecosystem, its uniqueness and global significance as a natural evolution experiment and the single largest reservoir of unfrozen freshwater, the microbial communities in the lake are just beginning to be characterized.

Historically, culture-dependent efforts focused on sewage runoff monitoring, with one amplicon effort using a taxonomically targeted approach [6]. Culture-independent efforts have explored oil seeps [7, 8], sediments, invertebrate gut [9], diatom-associated [10] and pelagic communities, with the latter on Sanger [11, 12] and 454 [13–15] platforms. Kurilkina and colleagues sampled a one station depth profile in September and June [14]. In both seasons, the main taxonomic groups included Proteobacteria, Actinobacteria, Bacteroidetes, Firmicutes, Chloroflexi, Acidobacteria, Cyanobacteria, while Caldiserica was substantially represented in September only.

Recent culture-independent studies emphasized analyses of metagenome-assembled genomes (MAGs), where contigs assembled from metagenomic shotgun sequences are binned into draft genomes. While these approaches offer additional insights into genes and function, they have been limited in scope to 1-5 samples per study. This lack of replication inherently limits the ability to connect genomic data with environmental covariates, as well as investigate co-occurrence patterns between taxa.

An amplicon-based survey with good spatial coverage was recently reported by Mikhailov *et al.* (2019), where the authors examined the possible biotic and abiotic influences on prokaryotic and eukaryotic communities. The survey data were from a June cruise, thus capturing the non-stratified homothermic period between spring and summer phytoplankton blooms. The study found correlation between overall community structure and environmental variables in multivariate space, as well as a few interesting cross-kingdom co-occurrences.

Here we present a comprehensive report of microbial communities during summer stratification in Lake Baikal. Our survey and analyses of microbial plankton span all three basins, from open waters to shallow bays, and multiple depths, including above and below thermocline, down to 500 m. We reveal major community composition trends that correlate with continuous environmental gradients in a spatial context. Multiple linear regression and model averaging reveals which specific environmental covariates are associated with the two most fundamental measures of diversity: richness and evenness.

We then identify strongest environmental covariates to community composition in a multivariate framework. Finally, we use co-occurrence network analyses to reveal clusters of OTUs with contrasting environmental associations and identify clades and taxa most responsible for maintaining the structure of each network cluster. We conclude by suggesting that temperature may be the main driver of the microbial community changes in Baikal in response to global environmental change.

## MATERIALS AND METHODS

Our survey was guided by the recorded natural history of Baikal [3, 16, 17] that divides the lake into eight distinct regions (Fig. 1a, S1). Among them, are the open Baikal basins, Chivyrkuy Bay, Proval Bay, and Selenga river plume. Chivyrkuy Bay was sampled extensively to capture transition from the shallow innermost bay (9 m depth) to open waters. Proval Bay and Selenga plume stations represented the two most eutrophic areas in Baikal. In total, we collected samples from 24 stations, of which 10 were sampled at various depths for a total of 46 samples. Temperature and dissolved oxygen profiles were measured with a YSI Instruments sonde (YSI, Inc., Yellow Springs, OH, USA). Whole water was taken using a Van Dorn sampler (Wildco, Inc., Yulee, Florida, USA), and 5 L were filtered onto 3 μm and 0.22 μm mixed nitrocellulose acetate membranes (EMD Millipore, Billerica, MA, USA) and stored at −20°C in RNAlater (Life Technologies, Grand Island, NY, USA) to capture particle-attached (3 μm filter) and free-living (0.22 μm filter) fractions of microorganisms. Genomic DNA was extracted using the Mo-Bio PowerSoil Kit (Mo-Bio Laboratories, Carlsbad, CA), following manufacturer’s protocol. Then, the V4 region of the 16S rRNA gene was amplified using 515F–806R dual-index primers sequenced on a MiSeq platform (250PE), as described previously [18]. Raw sequences and metadata needed to reproduce our results are available in GenBank under BioProject PRJNA530071.

**Figure 1:**
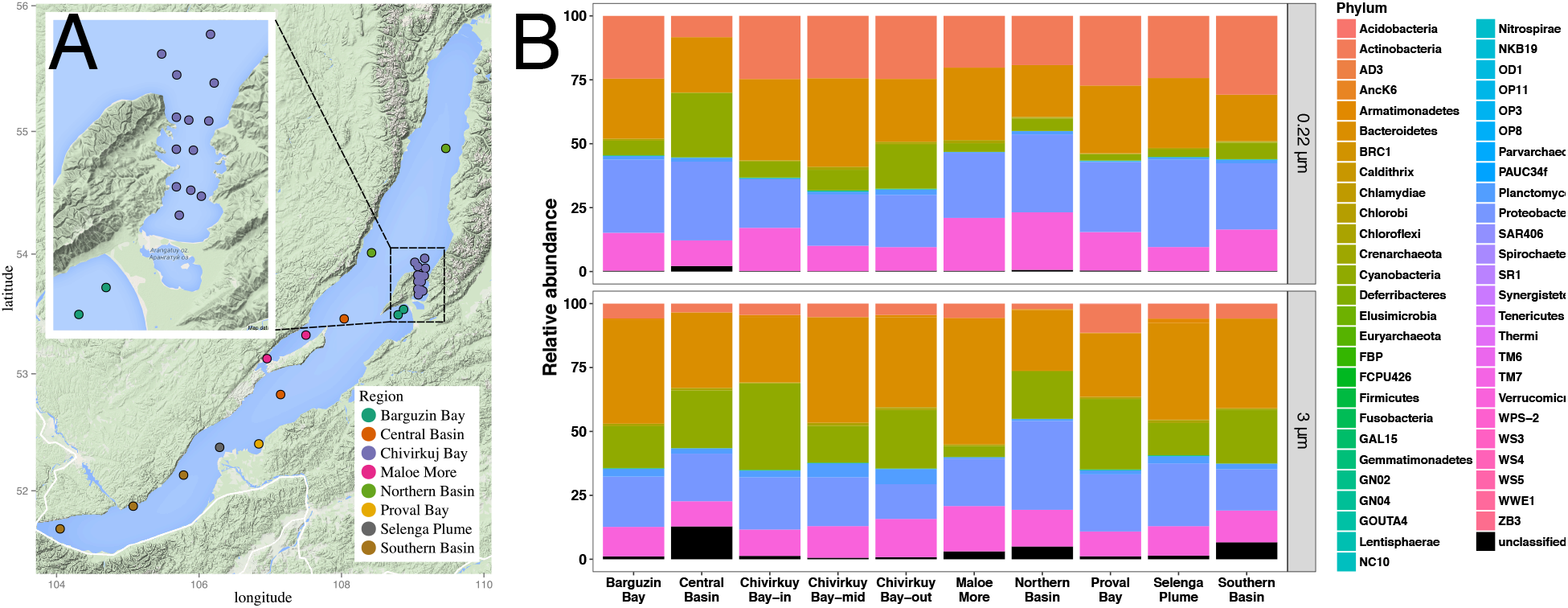
(A) Lake Baikal sampling locations. Stations in outside of Chivyrkuy Bay, Proval Bay, and Selenga Plume were sampled at multiple depths. (B) Taxonomic composition of the free-living (0.22 μm, top) and particle-attached (3 μm, bottom) size fractions across surface samples. While both fractions were mostly dominated by common freshwater phyla, Actinobacteria were enriched in the 3 μm fraction.

Sequences were processed generally following the *de novo* clustering mothur SOP. Statistical analyses were performed in the R (3.2.2) environment, unless noted otherwise. Multiple regression search and model selection were done with the glmulti package [19]. Distance matrices, ordinations, and correlations with environmental variables were calculated with vegan [20] and phyloseq [21] packages. We constructed the co-occurrence network using sparCC [22], following recommended best practices by Berry and Widder [23]. Next, we identified network modules with the optimum modularity algorithm in the iGraph package [24]. We followed the WGCNA package workflow [25] to generate eigenvectors for each module and correlate the first eigenvector (first principal component, PC1) with environmental variables to generate plots and a heatmap (Fig. 5).

## RESULTS AND DISCUSSION

### Taxonomic diversity

At a depth of 28059 sequences per sample, we detected 38457 OTUs in the combined 3.0 μm and the 0.22 μm fractions. Non-singletons (6099 in 3.0 μm and 3346 in 0.22 μm) were classified into 61 phyla, dominated by typical freshwater Actinobacteria, Bacteroidetes, Cyanobacteria, Proteobacteria and Verrucomicrobia (Fig. 1).

Microbial community composition differed from some previous studies of Lake Baikal, which explored pelagic communities [11–15]. In a recent effort, Kurilkina et al. [14] sampled one station depth profile in September and June to reveal the main taxonomic groups as Proteobacteria, Actinobacteria, Bacteroidetes, Firmicutes, Chloroflexi, Acidobacteria, and Cyanobacteria in both seasons, and Caldiserica in September only. Our study did not detect Caldiserica and revealed only a very minor presence of Chloroflexi, while showing a substantial Verrucomicrobia presence (absent in Kurilkina *et al.*) in every sampled region (Fig. 1). Interestingly, in the most recent amplicon survey of the lake [15], Mikhailov and colleagues report Verrucomicrobia relative abundance up to 59% in integrated water column samples taken across 0-25 m. The differences may be sensitive to the choice of amplicon primer sets, as Kurilkina *et al.*, Mikhailov *et al.*, and present study used three highly cited but ultimately different sets.

Our study revealed compositional differences between the 0.22 μm (free-living) and 3 μm (mostly particle-attached) fractions. Actinobacteria were significantly enriched in the 0.22 μm fraction (Table S1), similar to results from lakes in Michigan, USA [26]. Proteobacteria were only marginally more prevalent in the 0.22 μm fraction. We also found enrichment of Bacteroidetes and Cyanobacteria on the 3 μm fraction, the latter owing to filamentous taxa. Indeed, the top three most differentially abundant taxa identified with permutation-based analyses [27] were all classified as the Nostocales genus *Dolichospermum* and contributed up to 90% of Cyanobacteria in the 3 μm fraction. The high abundance of a nitrogen-fixing cyanobacterium confirms the significance of N limitation in the lake (O’Donnell et al. 2017).

Free-living (0.22 μm) surface samples in the open Baikal were dominated by betaproteobacterium *Limnohabitans* sp. Littoral zones were more variable, represented by multiple OTUs classified as *Limnohabitans*, *Synechococcus* and *Actinobacteria* acI. Surface samples of the shallow Proval Bay and the Selenga river plume represented the extreme end of the eutrophic gradient and were dominated by Actinobacteria acI and Verrucomicrobia *Chthoniobacter*. A representative from this genus is reportedly incapable of growth on amino acids or organic acids other than pyruvate [28]. Indeed, organic acids are among the most oxidized organic compounds that can be seen in a lake and would be found in areas where labile carbon is exhausted. *Chtinobacter*, therefore was found where expected - in eutrophic areas with ample labile carbon. Together with Bacteroidetes *Sediminibacterium* and *Cytophagia*, and Betaproteobacteria *Limonhabitans* and *Polynucleobacter*, the six OTUs make up ~46% of each Proval Bay and Selenga River plume’s relative abundance. The deepest samples in this study, collected at 500 m and 300 m, were dominated by Actinobacteria acI and acIV clades as well as ammonia oxidizing Group 1a Crenarchaeota *Nitrosopumilus*. The latter has been found in various environments. However, its type species is ubiquitous specifically in oligotrophic oceans, albeit in the surface layer and not at 500 and 300m [29].

### Community Richness and Diversity

Free-living community diversity showed opposing trends in the upper mixed layer (ML) and deeper waters (DW). Overall, richness and evenness strongly correlated with temperature in the ML and with depth in DW. For each sample, we calculated the effective number of species (ENS), a measure of diversity [30–32]. Hill [30] identified ENS as the number of equally-common species that yields a given value of a diversity index, such as the Shannon H’. Jost *et al.* [31] have argued that because ENS scales linearly with richness of equally-common species, it is the preferred metric for quantitative analyses (see Supplementary Methods). We show that the ENS increased with depth in DW (Fig. 2A) and was driven by an increase in evenness (Fig. 2C) with no accompanying trend in OTU richness (Fig. 2B). In the ML, ENS showed a modest increase at the surface in samples collected at 0 m (Fig. 2B). Unlike in DW, higher ML diversity was generated by higher OTU richness (Fig. 2B), while evenness remained unaffected. Mixed layer diversity was positively correlated with temperature (Fig. 2D), along with OTU richness (Fig. 2E). Deep water diversity and richness were marked by high overall variability, and richness showed a marginally significant increase in samples from cooler water. Evenness was not directly correlated with temperature in either layer (Fig. 2F). Furthermore, in mixed layer, richness had a greater response (slope) to temperature than diversity did, suggesting that samples at higher temperatures and shallower depths, while supporting the greatest richness, were dominated by a few successful OTUs. This could be due to variable and high resource conditions leading to coexistence of more taxa, including the persistence of rare taxa.

**Figure 2:**
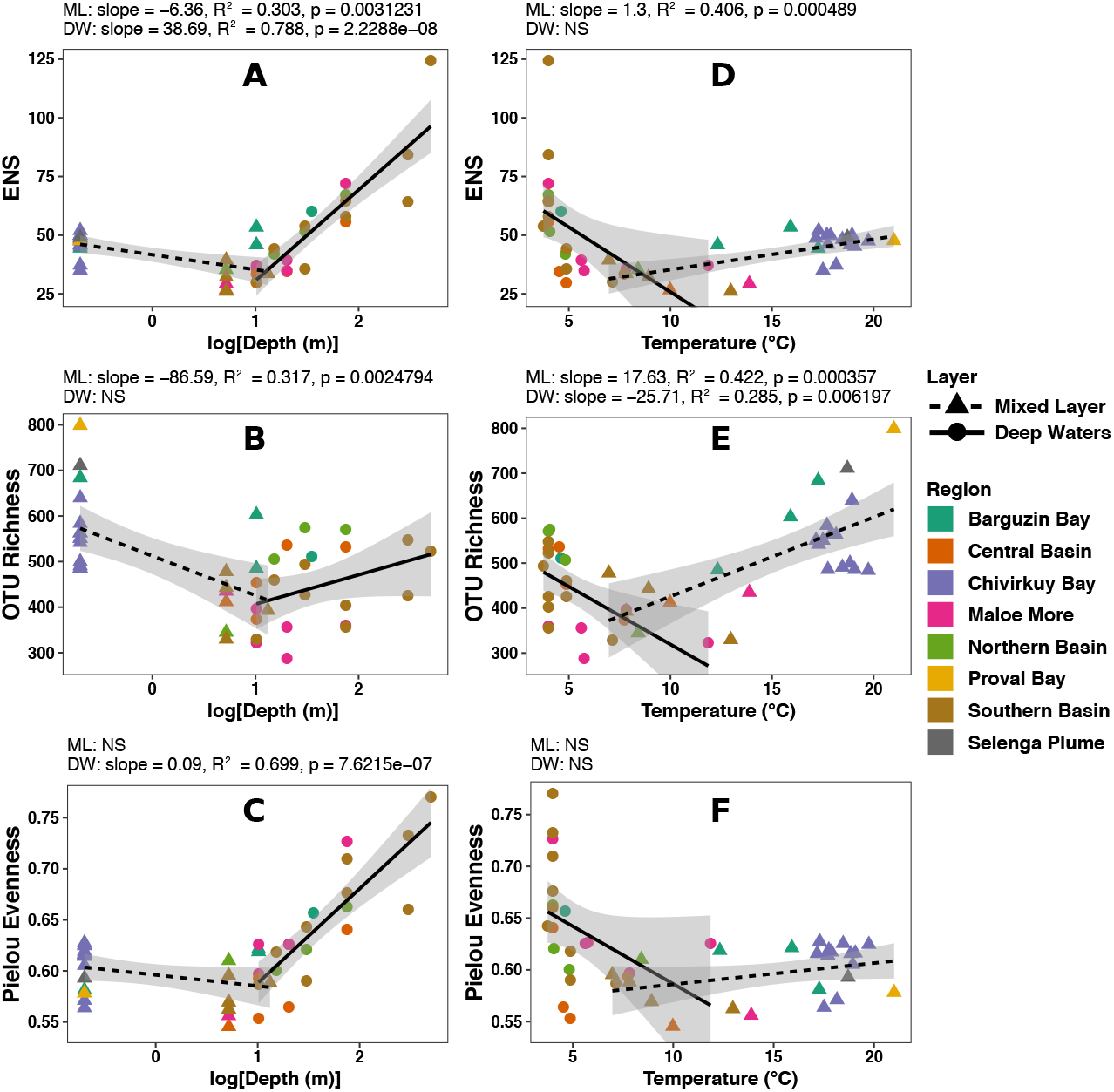
Diversity and evenness trends across depth and temperature in the upper mixed layer and deep waters. Effective Number of Species (ENS) was driven by richness in mixed layer and by evenness in deep waters. Effect size from Pearson correlation are above the panels.

Deep waters offer the opposite story. While richness had a significant but weak relationship with depth, ENS and Pielou evenness were very strongly positively correlated with depth (R^2^=0.79, p=2.2×10^−8^; R^2^=0.70, p=7.6×10^−7^), indicating that a substantial increase in diversity in the deep hypolimnion was not driven by richness but by an even community structure. These trends reveal an intriguing picture: warmer shallower part of mixed layer and deeper parts of the deep water layer both support more diverse (higher ENS) communities. But ML diversity, while rich in OTUs, is highly uneven, whereas the deep water communities achieve diversity by increasing evenness. The trends were similar for the 3 μm size fraction (Fig. S7). To better resolve underlying drivers that underpin these contrasting community features, we modeled ENS diversity, richness, and evenness with measured environmental variables using iterative multiple regression and model averaging [19].

Diversity trends in the ML were different, compared with deep waters. Diversity in the mixed layer correlated the most with temperature and total dissolved silica (TDS) (Fig 3A). These covariates were also significant for the OTU richness, while no measured environmental variable was a reliable predictor of ML evenness (Fig. 3B). Depth was not an important predictor for any ML community feature (Fig. 3C), possibly because depth variation was not large. These results highlighted the importance of temperature and silica in predicting the ML diversity, by elevating OTU richness. In the deep waters, total phosphorus (TP) and depth correlated with ENS (Fig. 3D). Furthermore, only TP was a significant predictor for the deep water OTU richness, while only depth was for evenness (Fig. 3E, F). Temperature, when controlling for other covariates, was not an important community feature predictor in deep water. The importance of temperature in ML and depth in DW was not unexpected, considering that the surveyed mixed layer spanned hundreds of kilometers across all three Baikal’s basins and multiple bays, where temperature ranged from a median 9.5**°**C in open waters to 21**°**C in Proval Bay (Fig. S1), while deep waters were characterized by a large range of depths from 10 to 500 m and a relatively constant temperature environment. Additionally, nutrients appeared to associate specifically with OTU richness. Dissolved silica, combined with light availability in ML, is known to promote diatom growth. Diatom exudates, acting as resources, could create additional niches, enabling coexistence of a greater number of OTUs. In deep waters, TP could promote OTU richness by supporting higher phosphorus demands often attributed to fast-growing copiotrophic organisms [33]. Logue and colleagues [34] found TP alone to be significantly correlated with OTU richness in a survey of Swedish lakes. Our results suggest that eutrophication leads primarily to an increase in the overall number of OTUs, but also to dominance of those with opportunistic lifestyles.

**Figure 3:**
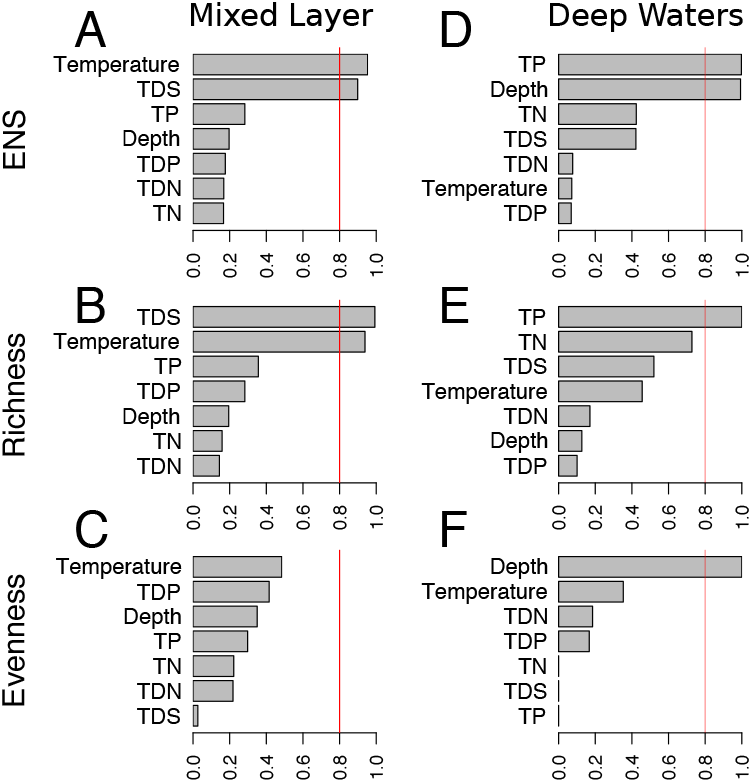
Model-averaged importance of environmental predictors for ENS diversity (top), OTU richness (middle) and Pielou Evenness (bottom) in the upper mixed layer (left) and the deeper waters (right). Tested predictors included total dissolved silica (TDS), total and dissolved phosphorus (TP, TDP), and total and dissolved nitrogen (TN, TDN).

### Multivariate trends

Ordination of the Bray-Curtis dissimilarity matrix of the 0.22 μm samples revealed clear grouping, which we designated into four significant clusters (Fig. 4A, permANOVA p<0.001). The first cluster (eutrophic) comprised samples from shallow and warm areas, specifically inner Chivyrkuy and Proval bays. The second cluster (transition) included samples from the Chivyrkuy Bay to open lake Baikal gradient, the main tributary Selenga River plume, and the two samples collected in Barguzin Bay – the deepest and most open of the three examined bays. The third cluster (open) contained the bulk of our samples collected in open waters that largely separated in ordination space along a depth gradient. The last cluster (deep) comprised the three deep water samples from 300 and 500 m. *Post hoc* pairwise comparisons of cluster centroids revealed significant differences between each cluster pair (FDR corrected p<0.05). Thus, the clusters broadly captured the different habitats of lake Baikal, separated by depth and the transitions between open lake waters and bays. Given the confounding effects of abiotic forcing and spatial autocorrelation, we used reciprocal causal modeling [35, 36] to test whether selection by abiotic environment or spatial dispersal/distance better explained community composition trends in mixed layer.

**Figure 4:**
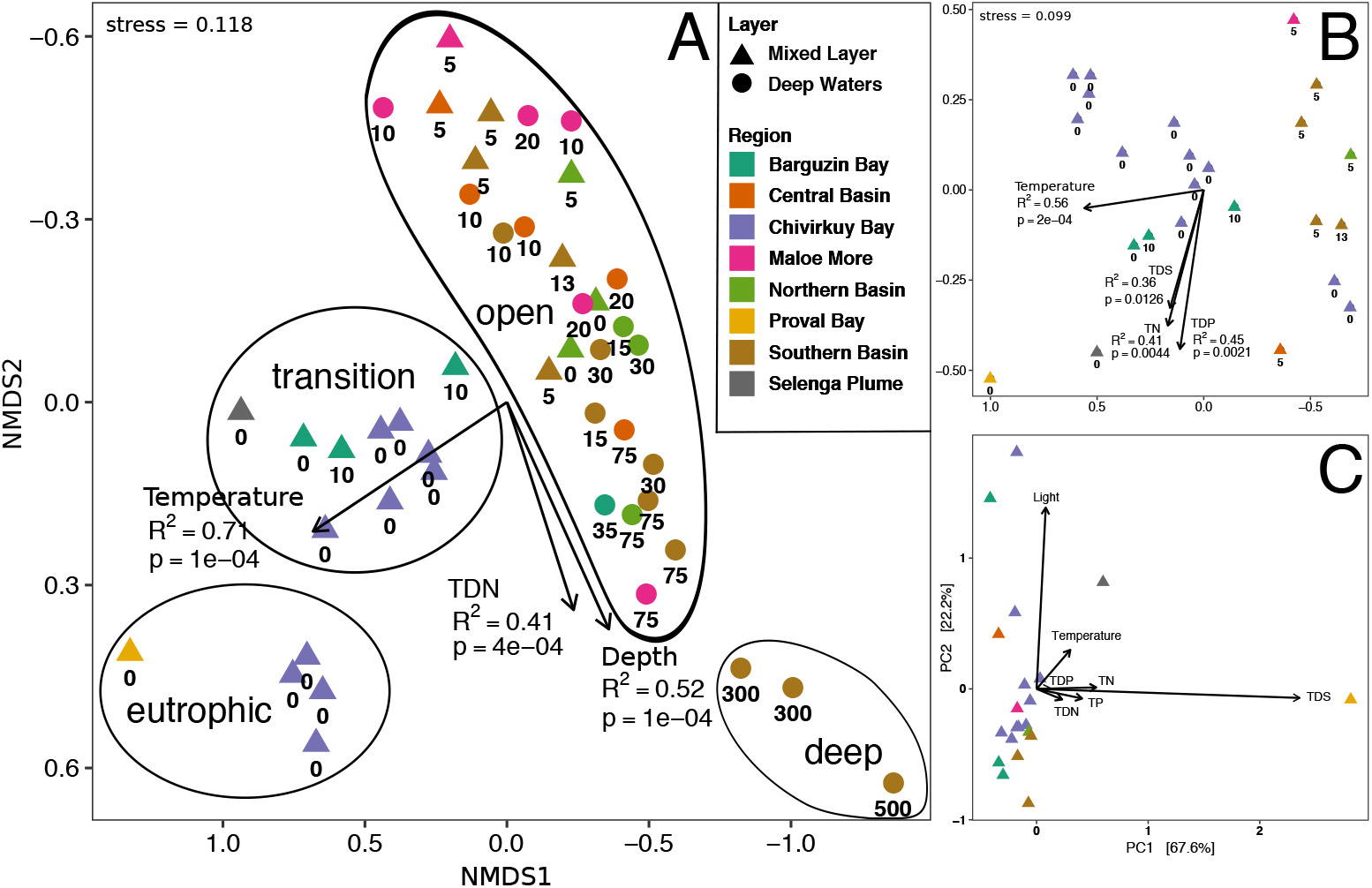
Ordination of the Bray-Curtis dissimilarity matrix for all 0.22 μm fraction samples (A) and the mixed layer only samples (B). (C) Principal component analysis of the mixed layer samples in environmental space using all measured covariates. In all panels, colors reflect major recognized regions of lake Baikal (15, 22). In (A) and (B), numbers below bubbles indicate sample depth (m), and arrows show correlation of environmental covariates with the layout of sample points in ordination space. Each arrow length = *R*^2^; p-values were obtained using permutations. In (A), ellipses were drawn to aid visualization.

Environmental conditions, and not the distance, played a dominant role in structuring free-living mixed layer communities. Reciprocal causal effects models test the hypotheses for significant correlation between the community dissimilarity matrix and each of the geographic distance and environmental distance matrices, while controlling for the other. We found that, when controlling for environment, geographic distance had no correlation with community dissimilarity. In contrast, when controlling for geographic distance, community structure showed a significant correlation with the environment (R^2^=0.33, p<1×10^−6^). Our results are consistent with other studies in freshwater [37, 38] and marine systems [39], which usually note stronger effects of environment on species composition, compared with dispersal, suggesting that dispersal is likely not limiting community assembly at this scale.

Temperature, depth, and total dissolved nitrogen (TDN) were the strongest environmental covariates with multivariate dissimilarity trends among free-living communities (Fig. 4). The temperature vector (Fig 4A, R^2^=0.71, p=1×10^−4^) indicated the greatest temperature variability along the open waters to bays gradient. We also confirmed the presence of the depth covariate among samples collected in the open waters (R^2^=0.52, p=1×10^−4^). Interestingly, open water community dissimilarities also revealed correlation with TDN in approximately the same direction as depth. Because depth can confound the effects of various environmental factors, we further considered an ordination of samples just from the mixed layer (Fig. 4B). Among those samples, temperature was still the strongest predictor of community dissimilarity (R^2^=0.56, p=2×10^−4^), and TDS was also significant (R^2^=0.36, p=0.012). Notably, these were also the only two model-averaged important predictors of OTU richness and ENS (Fig. 2A, B). However, TN and TDP were also significant predictors of community dissimilarity (Fig. 4B), although they did not show an association with alpha diversity metrics (Fig. 2A, B, C). As expected, depth was a not a significant predictor of community dissimilarity in the mixed layer. Lastly, we investigated whether the same environmental variables that predicted differences in microbial community structure also accounted for major abiotic differences between sampled sites. For this, we constructed a PCA of mixed layer samples using all measured environmental factors (Fig. 4C). The first two principal components captured 89.8% of the variation. TDS and light had the highest scores of all environmental variables. Furthermore, they were almost parallel to PC1 and PC2, respectively, suggesting that the two were responsible for explaining most of the measured abiotic differences between sampled ML sites. While TDS was a significant environmental covariate in ordination of biotic community structure, light was not (Fig. 4B). Similarly, light was not a significant predictor of biotic alpha diversity metrics either (Fig 2). TDP was the least important predictor of abiotic differences in ML (Fig. 4C); however, it was the second most significant predictor of biotic dissimilarity in ML (Fig. 4C). These features highlight that changes in microbial community structure were not correlated with simply the most variable environmental factors, and that significant associations of particular environmental factors warrant attention in further studies.

### OTU co-occurrence networks

To gain more insight into bacterial community structure in Lake Baikal and its dependence on the abiotic drivers and to identify OTUs that tend to co-occur, we constructed an OTU co-occurrence network. Co-occurrence networks enable data-guided identification of OTU assemblages and correlation of their abundance with continuous environmental variables. In contrast to ANOVA-based permutation procedures, networks do not carry a user bias of arbitrarily defining sample categories, which, even if chosen wisely, result in information loss.

Our network captured the bulk of Baikal’s OTUs. We constructed a co-occurrence network for OTUs present in at least 80% of samples (105 nodes and 819 edges, Fig. 5; see Supplementary Methods). Importantly, although 105 OTUs appeared to be a large reduction from the total >38,000 detected in Baikal, the network OTUs amounted to over >81% of cumulative relative abundance of every sample in >85% of samples. The remainder of the samples had 63% ± 17% of relative abundance included in the networks. Environments that were better sampled had more OTUs in the networks. Lower coverage environments included the notable outliers collected at 500 m, 300 m in Southern Basin and at the surface of the eutrophic Proval Bay and Selenga River plume. Thus, with network analyses we used non-categorized continuous data to directly capture the vast majority of microbial community and environmental covariate data, revealing the dominant groups of co-occurring OTUs and their association with major environmental drivers.

**Figure 5:**
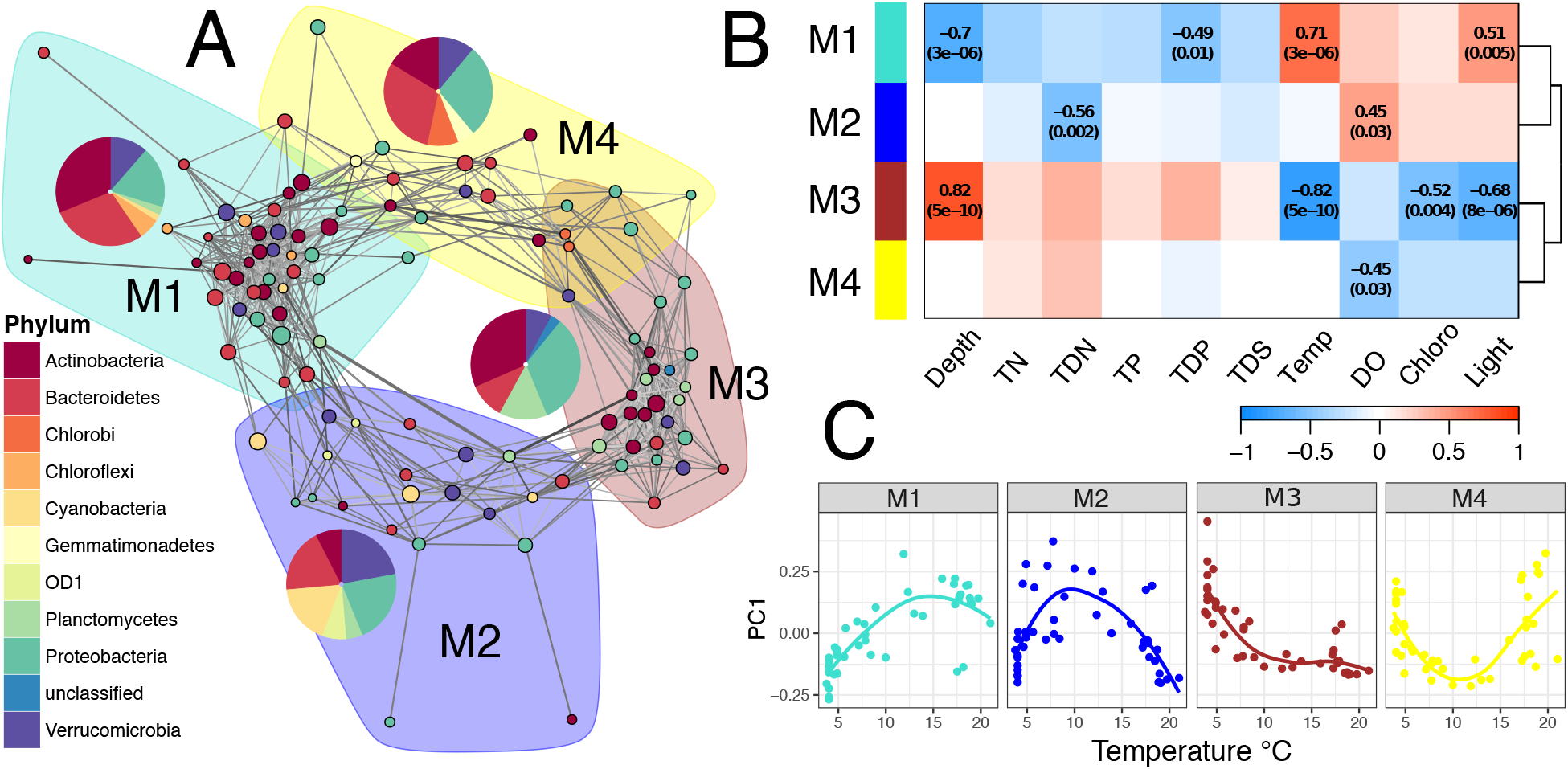
(A) Co-occurrence network of OTUs across all samples. Edge grayscale hue reflects pairwise correlation strength (darker edges show stronger correlations), and thickness indicates edge betweenness score. Modules were defined using maximum modularity optimization. For each module, the first principal component (eigenvector, PC1) of just that module’s constituent OTUs abundance matrix was used to summarize the dominant abundance trends across sampled sites. (B) PC1 trends are summarized in a heatmap, where numbers are Spearman correlation coefficients with p-values in parentheses below. Empty cells indicate non-significant results. (C) Example PC1 trends are shown with respect to temperature.

The resulting network exhibited the “small world” properties (Fig. S10). Small world networks are characterized by high connectivity between neighboring nodes and low connectivity between distant nodes [40], creating clusters or modules of consistently co-occurring OTUs. Within each module, OTUs with many connections (central nodes) are thought to reflect the processes that bring together module members. Small world networks may be more robust to perturbations but changes to the abundances of the central, well-connected OTUs may disproportionately affect the whole modules [41–43].

Using the optimal modularity approach [24, 44], we identified four modules of OTUs that tend to co-occur across sampled sites (Fig. 5A). Strong cohesion (see Supplementary Methods) within Module 1 (M1; clustering coefficient, CC_M1_=0.68; Table S1) and Module 3 (M3; CC_M3_=0.82) suggested common drivers for the consistently co-occurring OTUs. In contrast, low cohesion within Module 2 (M2; CC_M2_=0.54) and Module 4 (M4; CC_M4_=0.58) pointed to a looser internal structure.

Module eigenvectors [25] of M1 and M3 revealed opposing monotonic relationships with respect to environmental covariates (detailed introduction in Supplementary Methods). M1 showed a negative monotonic relationship with depth (Spearman ρ=-0.70, p=3×10^−6^) and a positive relationship with temperature (ρ=0.71, p=3×10^−6^). In contrast, M3 had a strong positive relationship with depth (ρ=0.82, p=5×10^−10^) and a negative relationship with temperature (ρ=-0.82, p=5×10^−10^). As expected, the direction of the opposing trends for the two clusters was reversed for light, owing to its inverse relationship with depth (Fig. 5B). Thus, we found that a large fraction of Baikal’s planktonic prokaryotes can be classified into one of the two cohesive clusters: mixed layer “warm” M1 and deeper waters “cold” M3 cluster.

Modules M2 and M4 (Fig. 5a) showed inverse opposing unimodal (non-monotonic) trends with depth, temperature and light (Fig. 5B, C). Weaker clustering within these modules suggested less cohesion, possibly due to the inclusion of taxa with weaker habitat preferences or terrestrial or riverine dispersal. M2 showed highest cumulative relative abundance at an intermediate temperature of about 10°C, with a strongly positive relationship with oxygen. In contrast, M4 showed a negative relationship with oxygen, abundance peaks at low and high temperature extremes and minimum abundance at approximately 11°C. Specifically, M4 was high in abundance in the immediate surface (0 m) and deep water, but with a sharp drop in abundance at 5 m depth.

### Phylogenetic signal in modules

Membership in the modules had a weakly significant phylogenetic signal (Fig. S11). This, combined with different habitat preferences among the modules, suggests that different taxonomic groups displayed preferential environmental associations, thus supporting the importance of niche differences and phylogenetic niche conservatism.

M1 and M4 had a high percentage of *Bacteroidetes*, known as opportunistic degraders of high molecular weight organic matter, such as proteins and carbohydrates, with genomes containing numerous carbohydrate-active enzymes covering a large spectrum of substrates from plant, algal and animal origin [45]. In aquatic systems, *Bacteroidetes* have been noted to follow pulses of organic matter inputs and cyanobacterial blooms [46]. This further casts M1 as a warm water ML module and M4 as the loose product of sediment and terrestrial input.

M1 is the only module to contain members (three OTUs) of the Chloroflexi phylum. Although Chloroflexi have been found in diverse environments, including oxygenated (its type genus) and anoxic [47] hot springs, the CL500-11 clade – a subclass of the deep-ocean SAR202 clade – has been suggested as characteristic of oxygenated hypolimnia in deep lakes, including Crater Lake [48], Lake Biwa [49] and the Laurentian Great Lakes [50]. However, the three Chloroflexi OTUs detected in our Lake Baikal network were from the Chloroflexi class (two OTUs) and Roseiflexales order (one OTU), and, as part of M1, appear to prefer warmer and shallower environments in the lake.

The cold deep-water cluster M3 had the highest relative abundance of OTUs in unclassified phyla. While the module also contained characteristically terrestrial Gammaproteobacteria *Pseudomonas* and *Acinetobacter*, their low connectivity suggested they are unlikely to play a central role in M3 processes.

M2 is the only module with Cyanobacteria; the module’s three most abundant OTUs – indeed, second and fourth most abundant in the entire >38,000 OTU dataset – classified as *Synechococcus*. Cumulative relative abundance of *Synechococcus* OTUs peaked at ~15 m, reflecting the approximate location of deep chlorophyll maximum at stratified stations. To further understand internal structure of the four modules, we focused on central OTUs that played important roles in creating the module structure.

### Central OTUs in each module reflect module ecology

The meaning of a node (in our case OTU) position in the network is the subject of much discussion. A central OTU has a significant positive correlation with a large number of that cluster’s OTUs. We offer one abiotic and one biotic interpretation, resulting from two explanations for the presence of network clusters in the first place. First, clusters of OTUs that correlate with important environmental covariates, like temperature, could reflect abiotic habitat filtering. In this case, OTUs in modules with higher clustering coefficients would more consistently reflect environmental conditions associated with that module. For example, every OTU in M3, which has the highest clustering coefficient (Table S1), shows clear negative relationship with temperature (Fig. S12). A second explanation requires an assumption that co-occurrences reflect biotic species interactions. Then, a central OTU could be hypothesized to directly increase the abundance of other OTUs, e.g., by synthesizing or otherwise making available a limiting resource. It is important to point out that the two explanations are not mutually exclusive and could both be parts of ecological mechanisms that give rise to observed clusters of co-occurring microbial species.

The most globally central (most connected) OTUs belong to the most populous module with the greatest number of connections (M1). The most central OTU in the network is OTU137, classified as an autotrophic methylotroph LD19, which was previously reported as a summer-fall bloomer in Lakes Michigan and Muskegon, MI [51]. Although not very abundant in our dataset, its high centrality suggests it is indicative of an ecological process that is at least partially responsible for supporting other OTUs in the ML.

OTU001 (*Limnohabitans*) – also in M1 – has a high overall abundance and the network position that straddles the balance between centrality in M1 and connectedness to M2. Its high relative abundance and ubiquity across the lake suggest important roles in lake ecology. Connectedness to two modules could reflect this ubiquity and further indicate an influence of OTU001 on OTUs that occupy both niches. More broadly, *Limnohabitans* genus has many ecotypes. Its fine-scale phylogeny and functioning, shown experimentally with isolates [52], revealed diverse lifestyles. In fact, three OTUs out of 105 in our network were classified as *Limnohabitans*, and one of the other OTUs (OTU104) was in the hypolimnetic M3. Separation of different *Limnohabitans* OTUs into M1 and M3 with opposing spatial occurrences and environmental preferences suggests they are indeed ecotypes [53] with distinct functions. Future studies can experimentally assess *Limnohabitans* ecotype responses to temperature and other environmental drivers.

The most connected OTU in the hypolimnetic M3 was Planctomycetes OTU022, classified into the Phycisphaerales order. Little is known about the ecological role of Planctomycetes in aquatic environments [46]. The order is known for its distinct visual appearance, and a recent report of a 3D reconstruction of Phycisphaerales cellular membrane revealed characteristic deep invaginations in the cellular envelope [54]. It is possible the increased surface area of Phycisphaerales becomes useful in the oligotrophic hypolimnion of Baikal. In contrast to the most central but not abundant OTU in M1, OTU022 is also the most abundant in M3, which is the most tightly clustered module in the network (Table S1). These data suggest Phycisphaerales may substantially contribute to nutrient cycling below the thermocline with direct impacts for the rest of M3.

The most recent amplicon survey of Baikal included co-occurrence network analyses of prokaryotes from the photic layer [15], calculated across samples collected at integrated depths from 0 to 25 m. Authors reported a similar phylogenetic makeup in a network that was characterized by predominantly positive interactions. Our sampling plan comprised separate samples from discrete depths in both the photic and abyssal layers, making it possible to resolve network clusters associated with the different conditions within the water column.

In summary, our survey of Lake Baikal’s bacterioplankton revealed that temperature, along with nutrients, are the major drivers of the microbial community composition and diversity, consequently, the anthropogenic changes in these factors would likely significantly alter planktonic microbial communities. The OTU co-occurrence network analysis identified two major clusters, associated with the upper mixed layer and deep waters, with a detectable phylogenetic signal in each cluster composition. The “small world” properties of the network suggest that the communities may be resilient to perturbations but the changes in abundances of the central OTUs may result in network rearrangement and substantial community shifts. Further studies, including controlled field and laboratory experiments, are needed to experimentally test the effects of different temperatures and other abiotic factors on the structure and diversity of microbial communities in Lake Baikal.

## Conflict of Interest Statement

The authors declare that the research was conducted in the absence of any commercial or financial relationships that could be construed as a potential conflict of interest.

## ACKNOWLEDGMENTS

We thank Ted Ozersky for valuable discussions, Lev Yampolsky for help with sampling plan design, and Pam Woodruff and Allyson Hutchens for assistance in the laboratory with nutrient concentration measurements. We are grateful to the crew of the R.V. *Treskov* of the Limnological Museum in Listvyanka and field technicians Elena Pislegina and Alexander Pislegin at the Institute of Experimental Biology, Irkutsk State University for facilitating fieldwork. We thank Ashley Shade and Jim Tiedje for comments on the manuscript. This study was funded by National Science Foundation (NSF) Dimensions of Biodiversity grant DEB-1136710.

## Notes

### Competing Interest Statement

The authors have declared no competing interest.

### Summary of Updates

Edited manuscript for journal submission.

