## Supplement for "Environmental drivers define contrasting microbial habitats, diversity, and community structure in Lake Baikal, Siberia"

### SUPPLEMENTARY MATERIALS

#### MATERIALS AND METHODS

*Sampling and environmental contextual data*

The survey was guided by the recorded natural history of Baikal [1, 2] that divides the lake into eight distinct regions. We collected samples from 24 spatial locations, where 10 of those locations were sampled at various depths for a grand total of 46 samples across the lake (Fig. S1). The cruise took place on board the R.V. Treskov on August 3-17, 2013.

Temperature and dissolved oxygen profiles were measured with a YSI Instruments sonde (YSI, Inc., Mod4 Springs, OH, USA). Light profiles were measured using Walz model US- SQS/L model Li-185B light probe (Heinz Walz GmbH, Germany), connected to a Li-Cor Quantum photometer (LI-COR Biosciences, Lincoln, NE, USA).

Whole water was taken using a Van Dorn closing bottle (Wildco, Inc., Yulee, Florida, USA). From each sample, (a) 1L was filtered onto a GF/F glass fiber filter (GE Healthcare Bio-Sciences, Pittsburgh, PA, USA) for Chl a measurements and frozen at −20°C; (b) 5 L were sequentially filtered onto 3 μm and 0.22 μm mixed nitrocellulose acetate membranes (EMD Millipore, Billerica, MA, USA) and stored at −20°C in RNAlater (Life Technologies, Grand Island, NY, USA) to capture particle-attached and free-living fractions of microorganisms [3]; (c) 50 mL were frozen at −20°C for total nutrient analysis; and (e) 15 mL were filtered through a 0.22 μm mixed nitrocellulose acetate membrane (EMD Millipore) and frozen at −20°C for dissolved nutrient analysis.

Samples were transported to the USA in insulated containers cooled with liquid nitrogen. In the USA, samples for molecular analyses were stored at −80°C and samples for nutrient analyses were stored at −20°C.

*Temperature and light profile modeling*

Temperature values at collection sites were estimated from YSI instruments sonde profiles. YSI sonde profiled temperature at each station down to between 40 m to 50 m. Five temperature values per second were recorded on the downcast (lowering approximately 1 m per second) and used for modeling. For each station, high-order polynomial functions were fit to the data, and point estimates were used to infer temperature values at exact depths used for water sample collection (Fig. S2). For sites samples collected at 75 m, 300 m and 500 m, temperature was assigned a value of 4°C, based on literature values [4].

Light values at collection sites were estimated from profiles collected using the Li-Cor Quantum photometer (LI-COR Biosciences, Lincoln, NE, USA). Profiles were measured down to site bottom or maximum 20 m depth. The light extinction function $I_D = I_0e^{-kD}$, where $I_D$ = light intensity at depth, $I_0$ = light intensity at the surface, k = extinction coefficient (0.035 for pure water) and D = depth, was fit to the to the data to estimate light values at collection sites (Fig. S3).

*Nutrient Measurements*

Samples for nutrient analysis were thawed and digested with potassium persulfate at 120°C for 30 min. Total and dissolved nitrogen were measured colorimetrically on a Shimadzu UV-240-PC spectrophotometer (Shimadzu Scientific Instruments, Columbia, Maryland, USA) at 224 nm using the 2nd derivative method [5]. Total and dissolved phosphorus was measured using the orthophosphate method on a Lachat Instruments Quick Chem 8500 autoanalyzer (Lachat

Instruments, Loveland, Colorado, USA). Chl *a* filters were extracted with ethanol, and Chl *a* was determined fluorimetrically [6].

*DNA extraction and amplicon sequencing*

Genomic DNA was extracted using the Mo-Bio PowerSoil Kit (Mo-Bio Laboratories, Carlsbad, CA), following manufacturer's protocol. Then, the V4 region of the 16S rRNA gene was sequenced using dual-index primers as described previously [7]. After PCR amplification, the products were normalized and pooled. The pool was loaded on an Illumina MiSeq v2 flow cell and sequenced with a standard 500 cycle reagent kit for paired-end 250 bp reads (PE250). Base calls were done with Real Time Analysis software v1.18.54. Output of RTA was demultiplexed and converted to FastQ with Illumina Bcl2Fastq v1.8.4.

*Amplicon sequence processing*

The FastQ output files were processed using mothur, following general MiSeq protocol and the options below [7, 8]. Sequences were aligned to a full SILVA v.132 database. Chimeras were removed with UCHIME in mothur environment. The remaining sequences were classified with a naïve Bayesian RDP classifier [9] using the SILVA database. Sequences classified as Mitochondria, Chloroplasts and Eukaryota were removed, resulting in 197,738 unique sequences that were clustered into OTUs at 97% similarity. Consensus taxonomy for each OTU was determined following mothur protocol. Coverage for sequenced samples varied between 24348 and 76838 (Fig. S5, left), and was rarefied to the lowest coverage sample i.e. 24348 reads (Fig. S5, right).

*Multiple regression*

We used the brute force approach to multiple linear regression (exhaustive search, followed by model selection) to determine the relationship of community richness, diversity, and evenness with measured environmental variables. Richness was taken as the number of OTUs in each sample. This comparison was possible because our sampling effort was rarefied in mothur (see above). The effective number of species (ENS), represented diversity and was calculated using the Shannon diversity index H' [10, 11], as shown in Eq. 1.

$$ENS = e^{H'} \qquad \text{Eq. 1}$$

It was important to use ENS because, unlike diversity indices, such as Shannon and Simpson, ENS has a linear response to change in community diversity – a necessary property of a response variable in quantitative modeling [10, 11]. Statistical modeling and model selection were performed with the glmulti package in R [12].

*Multivariate statistical analyses*

Distance matrix. Ordination of the OTU abundance matrix was performed in the R environment (version 3.2.2) using vegan [13] and phyloseq [14] packages. First, we compared different techniques for calculating community dissimilarities. We used the Mantel test to reveal correlations between the Bray-Curtis, Jaccard, unweighted Unifrac, and weighted Unifrac distance matrices. Results showed greatest differences between presence-absence distance metrics and abundance-weighted distance metrics. However, within each of the two categories, the metrics were approximately interchangeable. Thus, we decided to use the Bray-Curtis metric for better compatibility with other studies. The high correlation of weighted Unifrac matrix indicated that phylogenetic information did not largely affect distance matrix results.

Test for dispersion (geographic distance) was done with reciprocal causal modeling using partial Mantel tests. The use of reciprocal partial mantel tests for reciprocal causal modeling is advocated in the literature by Cushman [15, 16], where he criticized simple Mantel tests for their Type I error rates, and proposed reciprocal partial mantel as a solution [15], later expanding to a more sophisticated "relative support" technique - also based on partial Mantel tests [16]. Rousset criticized all Mantel-based options in favor of mixed effects regression methods [17] because they are not robust when there is autocorrelation in the environment matrix. In a summary review, Cushman conceded that LME is indeed the best method for decoupling the effect of spatial dispersal and selection; however, that doesn't mean reciprocal partial mantel tests are inappropriate, especially if autocorrelation in data is weak [18]. Cushman pointed out that reciprocal partial mantel tests performed almost as well as mixed linear models if conclusions were based on $R^2$ effect sizes, and not just p-values. Fortunately, *our data does not have significant spatial autocorrelation with respect to environment* (Moran's I > 0.05). Thus, we proceeded using Cushman's method for its simplicity.

Geographic distance between sampled sites was calculated with the geosphere R package using the Vincenty Ellipsoid model. Mantel test was performed using the vegan R package. Because the geographic distance matrix was not normally distributed (Fig. S13), we used the rank-based Kendall test statistic option in the Mantel tests.

Ordination was done in the phyloseq package [14] and modified for visualization with a custom R script. NMDS plots are freely rotatable and scalable, and Fig. 4 panels a and b were thus adjusted for greater visual clarity.

Correlation with environmental variables was done with the envfit function in the vegan package, which correlates continuous environmental values with separation of points in ordination space. Significance was calculated by bootstrapping using 9999 permutations.

*Network construction and analyses*

Co-occurrence matrix and network construction. The OTU co-occurrence matrix was calculated for OTUs that were present in 80% of all samples (109 OTUs) with sparCC software [19], as recommended in best practices for co-occurrence network construction by Berry and Widder [20]. Significance values for correlations were bootstrapped, as described by sparCC authors, and then corrected for multiple testing using FDR manually in R with a custom script. Positive co-occurrences with adjusted $p < 10^{-5}$ were used for downstream analyses. The network was constructed, displayed and analyzed using the iGraph package [21] in the R environment. Two pairs of OTUs that were only connected to each other but not the rest of the network were removed, resulting in 105 OTUs in all subsequent analyses.

Comparison with simulated networks. We compared basic network statistics with null distributions of two types of simulated networks. First, we created 10,000 random Erdös-Rényi networks, which were undirected, without loops and used the gnm model with the same number of nodes (105) and edges (814) as our Baikal network. We also created 10,000 small-world Watts-Strogatz networks with 105 nodes and replacement probability p=0.05. Average path length and the clustering coefficient (transitivity) of the lake Baikal network were compared to distributions of simulated networks (Fig. S10). Two-tailed *p*-values for Baikal network statistics were calculated as the number of simulated observations greater than the absolute values of the Baikal network divided by the total number of simulations.

The network exhibited small world properties. Both overall CS (0.618) and the average path length (AP=2.62) were much higher than random Erdös-Rényi simulations (CC mean=0.150, p=0, n=10,000; AP mean=1.93, p=0, n=10,000; Fig. S10a) and higher than small-world Watz-Strogatz simulations (CC mean=0.514; p=0, n=10,000; AP mean=2.27, p=0, n=10,000; Fig. S10b). The basic network statistics are summarized in Table S1.

Small-world properties place greater topological importance on central nodes (nodes with many connections) and to a lesser extent bottlenecks in maintaining the network structure [22, 23]. Hubs can be defined as nodes with high eigenvector centrality, and bottlenecks as high betweenness nodes. Hubs are well-connected, in many cases to other well-connected nodes and are therefore considered topologically more central. In an ecological co-occurrence network, they have been hypothesized to mediate processes important to their neighbors [24]. For example, a central OTU may produce a common good limiting resource, like a vitamin.

Bottleneck nodes are positioned along a high number of shortest paths between pairs of other nodes. This is called high betweenness. Fig. 5 displays betweenness values for each edge as proportional to its width. In a small world network with few bridges between modules, bottlenecks are the only way one module can interact with another. Therefore, bottlenecks have been hypothesized to mediate feedbacks between ecologically meaningful OTU assemblages [24]. These features emphasize the strength of network methods to detect candidate keystone taxa not necessarily based on their large abundance but on how they affect other network players.

Community (module) detection. Network modules were identified using the optimal modularity algorithm [25] in the iGraph package. Optimal modularity calculates the arrangement and membership of clusters that gives the highest modularity score, as defined by Newman *et al* [23] and adopted by the iGraph authors.

Environmental trends of modules. To summarize OTU abundance trends across sampled sites, we used the first principal components of OTU abundance matrices (eigenOTUs [26]) for OTU members of each of the four modules. The resulting four eigenOTUs were used to assess correlation of modules with environmental factors, such as temperature, nutrients, oxygen, and chlorophyll levels. For example, module M1 has 39 member OTUs. They are together because, by definition, they have similar occurrence patterns across samples. But how can we summarize all OTU abundances in one vector? One option is to sum their relative abundances in each sample and get per sample cumulative values. However, this option is heavily biased towards the more abundant OTUs. Indeed, the difference between the most and least abundant OTUs in M1 is approximately two orders of magnitude. A better option is to use the widely practiced principal component analysis (PCA), which weighs OTUs by their power to explain variation in other OTUs. In PCA, the first principal component (PC1) is the best summary of OTU abundances in each sample. Thus, we ran an independent PCA for OTU members of each of the four modules and used PC1 for each module as a summary of its abundance across samples. The PC1s were correlated with measured environmental variables (Fig 5b, c).

Phylogenetic signal calculation. We first constructed a phylogenetic tree of OTUs by computing pairwise DNA distances using the K80 model and constructing a tree with the bionj algorithm [27]. Phylogenetic signal was detected using a modified phylo.signal.disc R script developed by Enrico Rezende (Universidad Autònoma de Barcelona). This function was used in several studies to reveal a phylogenetic signal in discrete traits [28–33]. It works with a tree of network OTUs and treats module assignment of each OTU as a character state with random transitions. The strongest possible phylogenetic signal in the OTU tree would be indicated by *num* - 1 transitions, where *num* is the number of modules. To quantitatively assess the phylogenetic signal, the script first uses maximum parsimony to generate a null model by calculating the minimum number of character transitions needed to create permuted character states of the OTUs for 10,000 simulations. Then, the actual number of transitions in the tree of Baikal OTUs with known module membership is compared with a distribution of transitions obtained from the permuted simulations. A p-value for a phylogenetic signal is calculated from comparing the true number of transitions with a null distribution of transitions obtained from the permuted simulations (Fig. S11).

#### SUPPLEMENTARY RESULTS

*Temperature, light and nutrient profiles*

Most stations were thermally stratified, while others exhibited weaker stratification because of the storm at the end of the cruise (Fig. S2). At the surface, temperature ranged from 7.0°C in northernmost SB to 21°C in Proval Bay with a median at 17.4°C. In the open waters, the median temperature was 9.5°C, consistent with expected values [1].

Light profiles with models are shown in Fig. S3, top. Light extinction coefficients (Fig. S3, bottom) were greatest at the single station in the eutrophic Proval Bay, high in the Selenga River plume and lowest in Maloe More. Central Baikal stations, sampled along the gradient from shallow bay to open lake conditions, showed variable values.

Nutrient profiles are shown in Fig. S4. Total nitrogen (TN) ranged from 3.0 μM in southern Maloe More Straight to 25.2 μM in Proval Bay with both an overall and open waters medians at 10.0 μM. TP ranged from 0.16 μM in northernmost South Baikal to 1.4 μM in Proval Bay with lakewide and open waters medians at 0.35 and 0.38 μM. Molar TN/TP ratios varied from 5.6 in southern Maloe More to 56.9 in southern Central Baikal; median TN/TP was 26.1. Chl a ranged from 0.66 μg L1 in mid-Chivirkuy Bay to 10.5 μg $L^{-1}$ in Proval Bay with the median at 2.0 μg μg $L^{-1}$. Nutrients increase, but with mild trends at depths above 80 m (not significant) with a visible increase at the deeper stations at 300 and 500 m. While relationships with TN, TP, DN, DP, DS and chlorophyll were all significant using all samples, only TN and chlorophyll remained significant with two high nutrient outlier stations (Proval Bay and Selenga Shallow) removed. The remaining positive relationship illustrated an expected correlation between nutrient load and primary productivity.

*Amplicon sequence processing*

see methods

*Multiple regression*

Trends on the particle-attached 3um fraction were similar to the free-living 0.22 um fraction (Fig. S6, S7)

*Multivariate statistical analyses*

Dissimilarity matrices based on phylogenetic distance metrics (unweighted and weighted unifrac) correlated very highly with phylogeny-free Jaccard and Bray-Curtis matrices (Fig. S8) and were concluded not to add significant additional information to modeling the multivariate community structure.

Ordination of the 3um fraction showed similar results to the 0.22um fraction (Fig. S9).

*Network construction and analyses*

see methods

#### SUPPLEMENTARY FIGURES

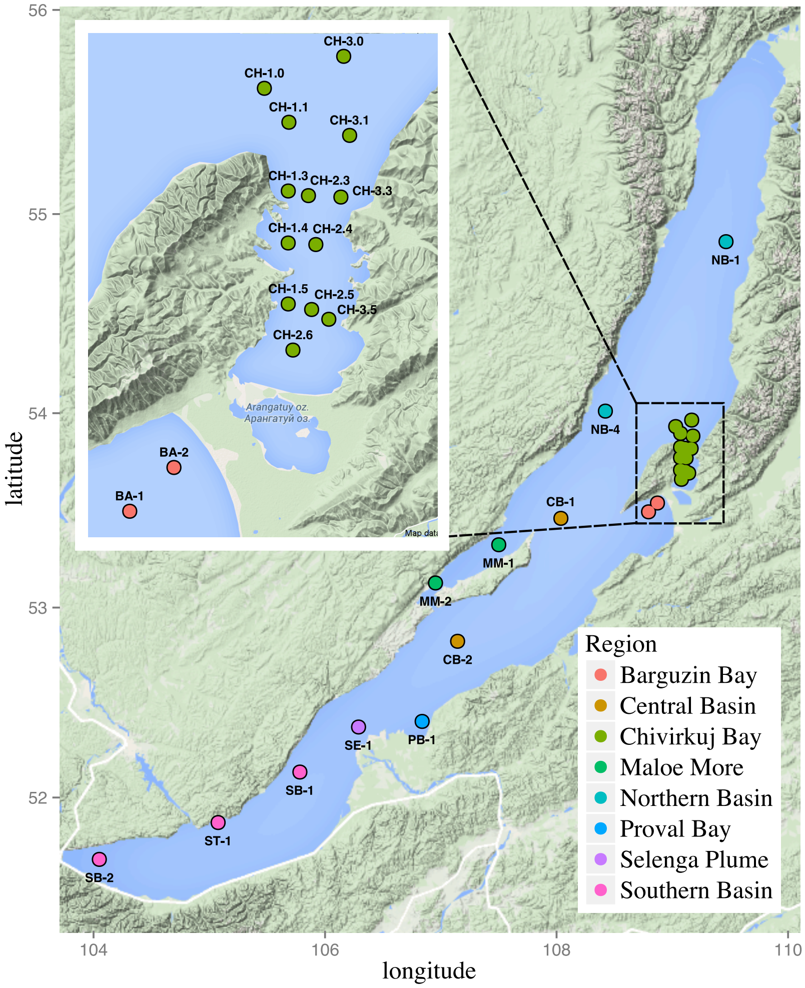


**Fig. S1:** Sampling sites at Lake Baikal. Colors reflect major recognized regions of lake Baikal [1, 4].

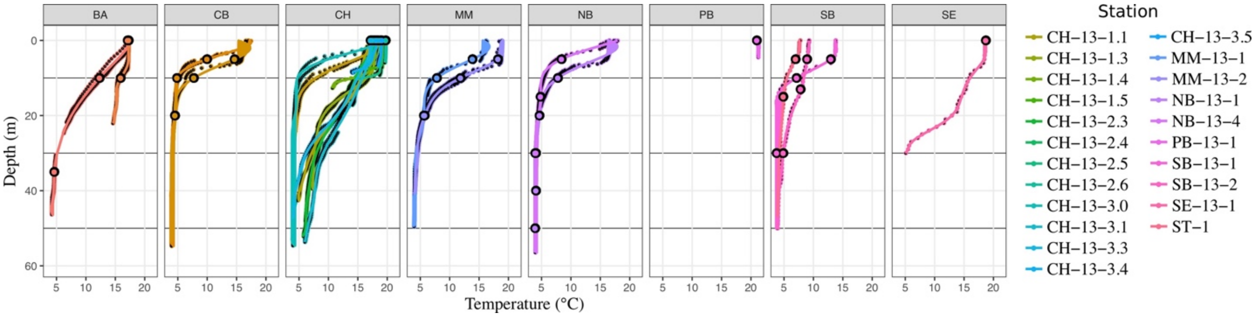


**Figure S2:** Temperature profiles and location of collected samples in the water column.

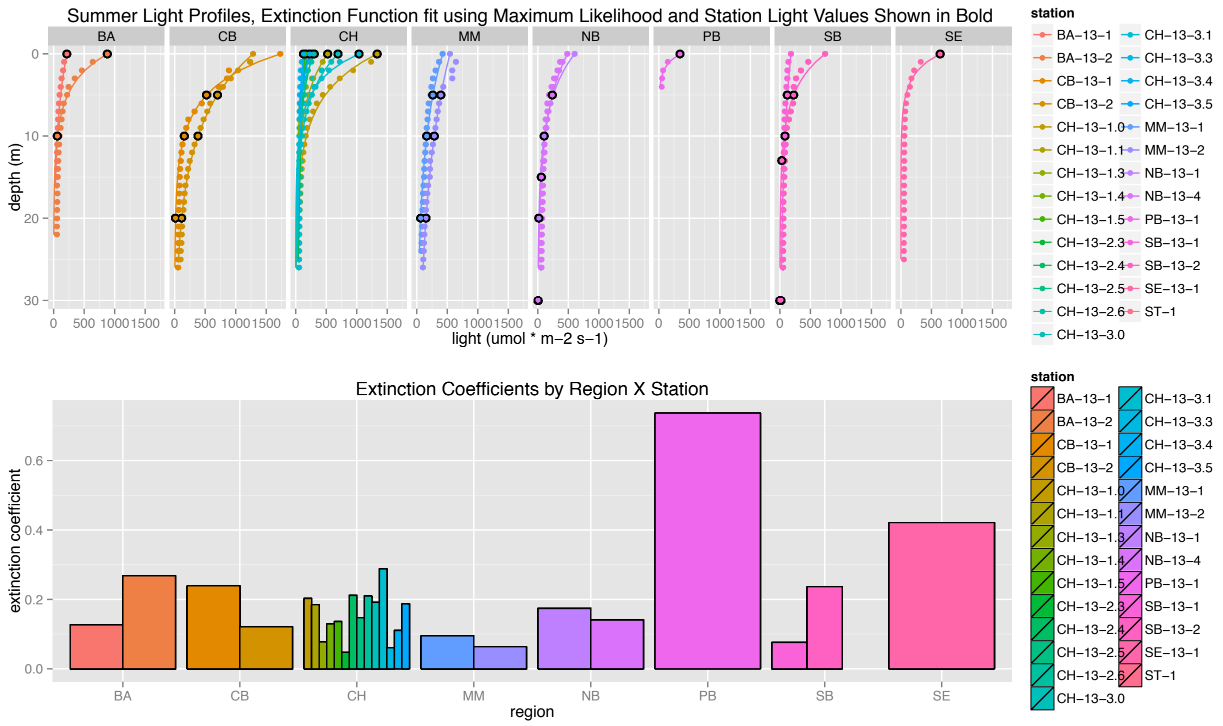


**Figure S3:** Light profiles, fitted models and extinction coefficients.

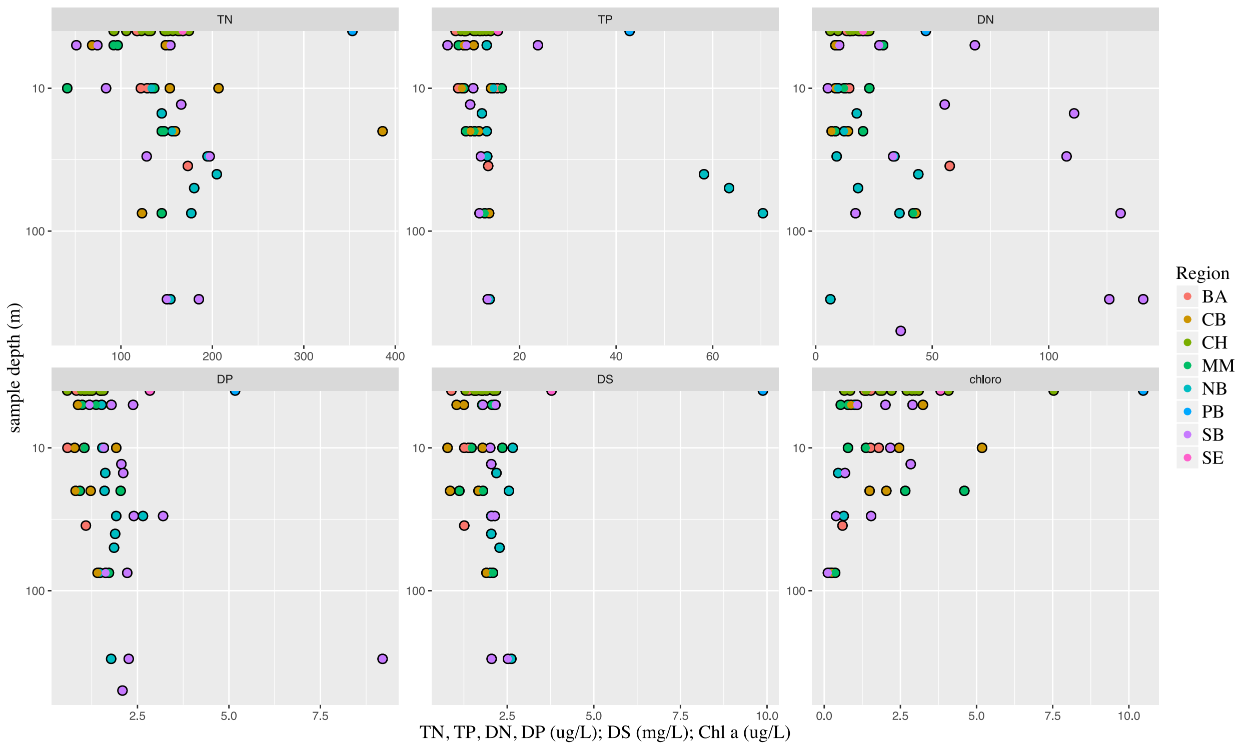


**Figure S4:** Nutrients show non-significant increase with depth.

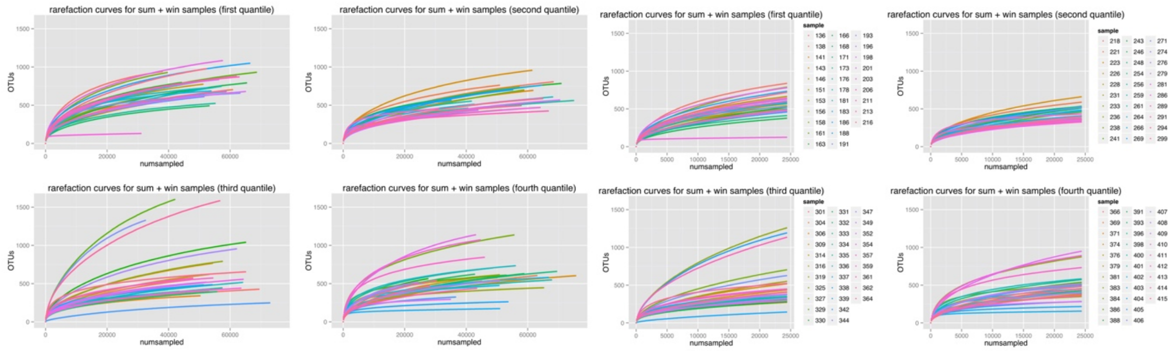

**Figure S5:** Rarefaction curves showing sampling depth per sample (left) and rarefied samples (right)

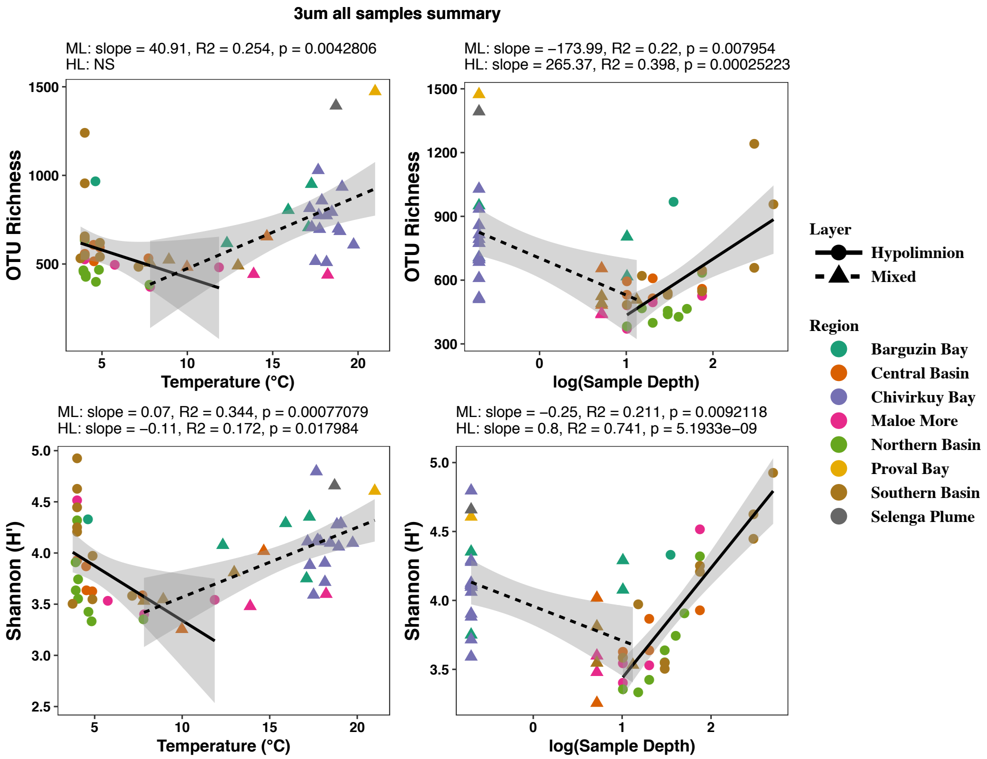


**Figure S6:** OTU richness and Shannon diversity on the 3 um fraction size, showing the same trends as the 0.22 um fraction.

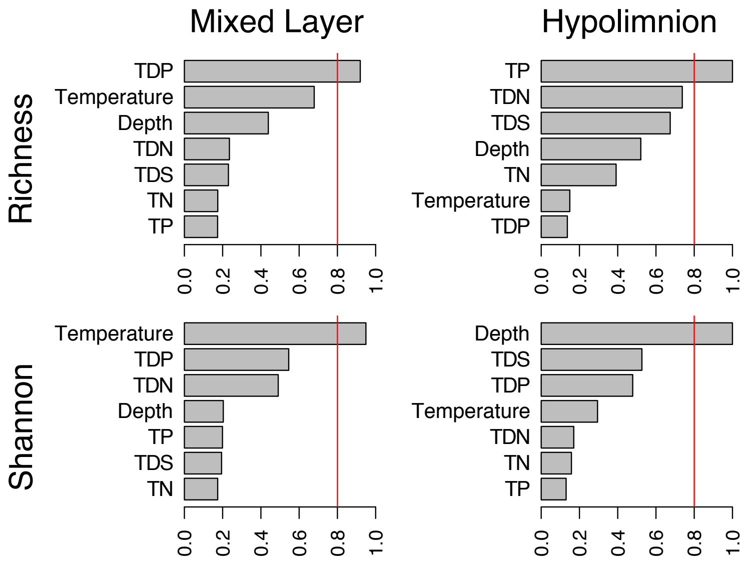


**Figure S7:** Model-averaged importance of environmental predictors for OTU richness (top) and Shannon diversity (bottom) in the mixed layer and the hypolimnion – on the 3 μm size fraction.

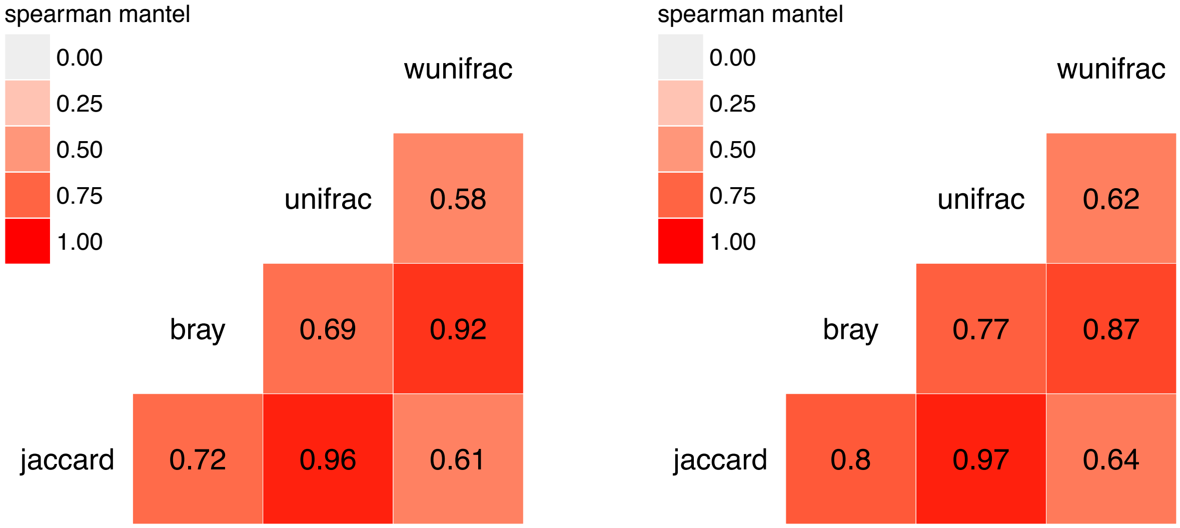


**Figure S8:** Pairwise Mantel test correlations between distance matrices based on phylogeny-free and phylogenetically-informed metrics for the 0.22 um (left) and 3um (right) size fractions. Unweighted distance calculators treat all OTUs equally, while weighted versions emphasize differences among the more abundant OTUs.

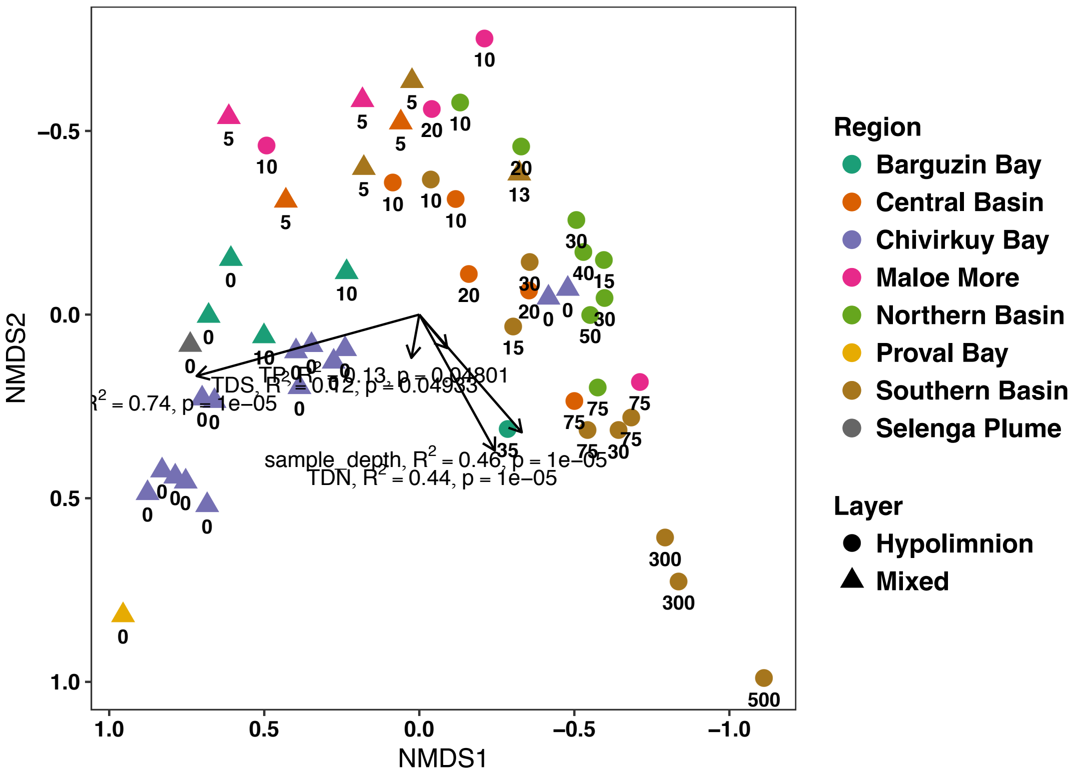


**Figure S9:** NMDS ordination of the particle-attached 3um fraction

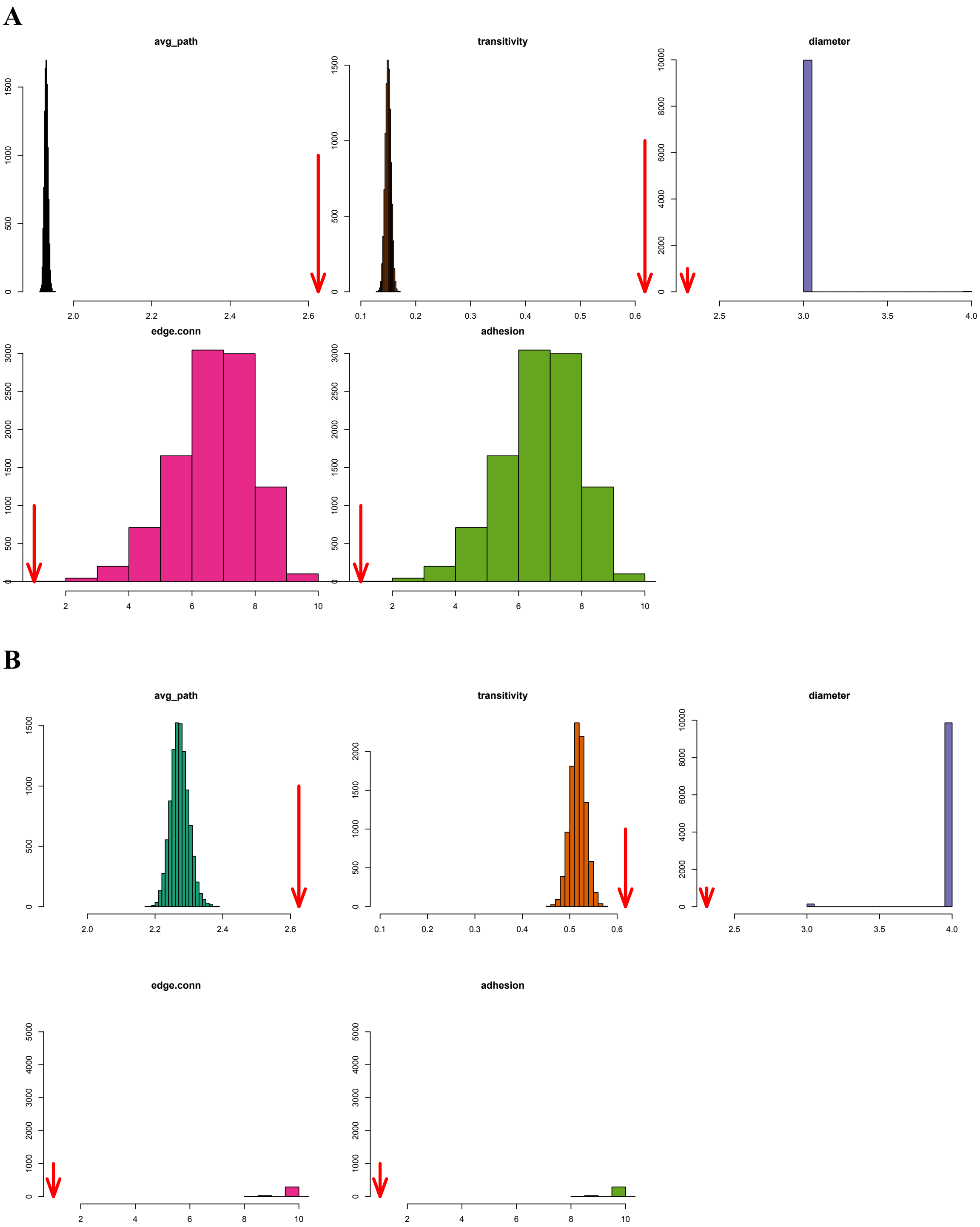


**Figure S10:** Network statistics comparison with simulated Erdos-Renyi (A) and Watts-Strogatz (B), run for 10,000 simulations each. Red arrows indicate statistics for Baikal networks.

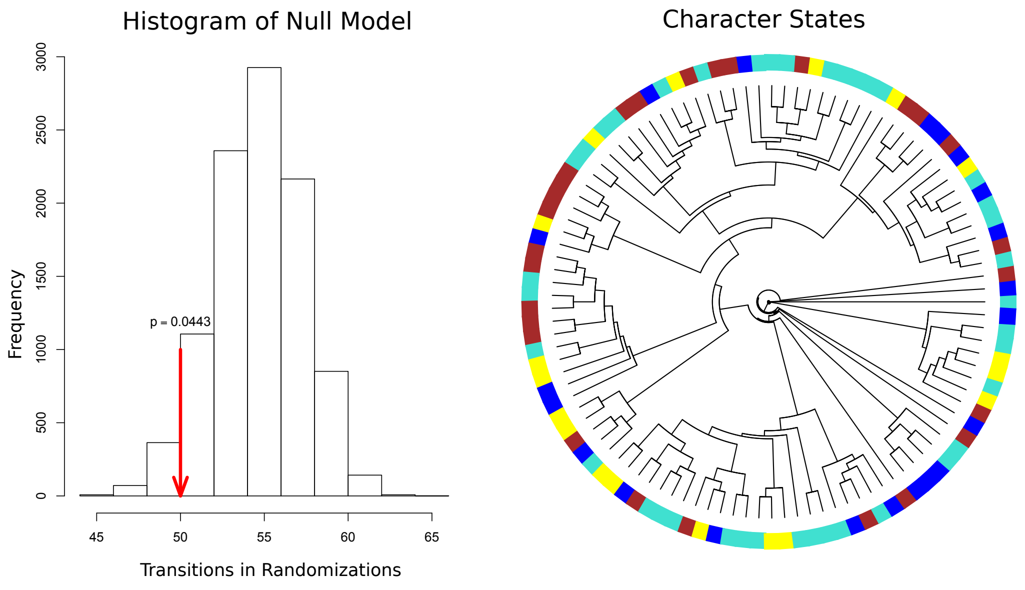


**Figure S11:** Phylogenetic signal for co-occurrence network modules as discrete character states for individual OTUs. Phylogenetic distribution was non-random across modules, possibly reflecting their ecological roles.

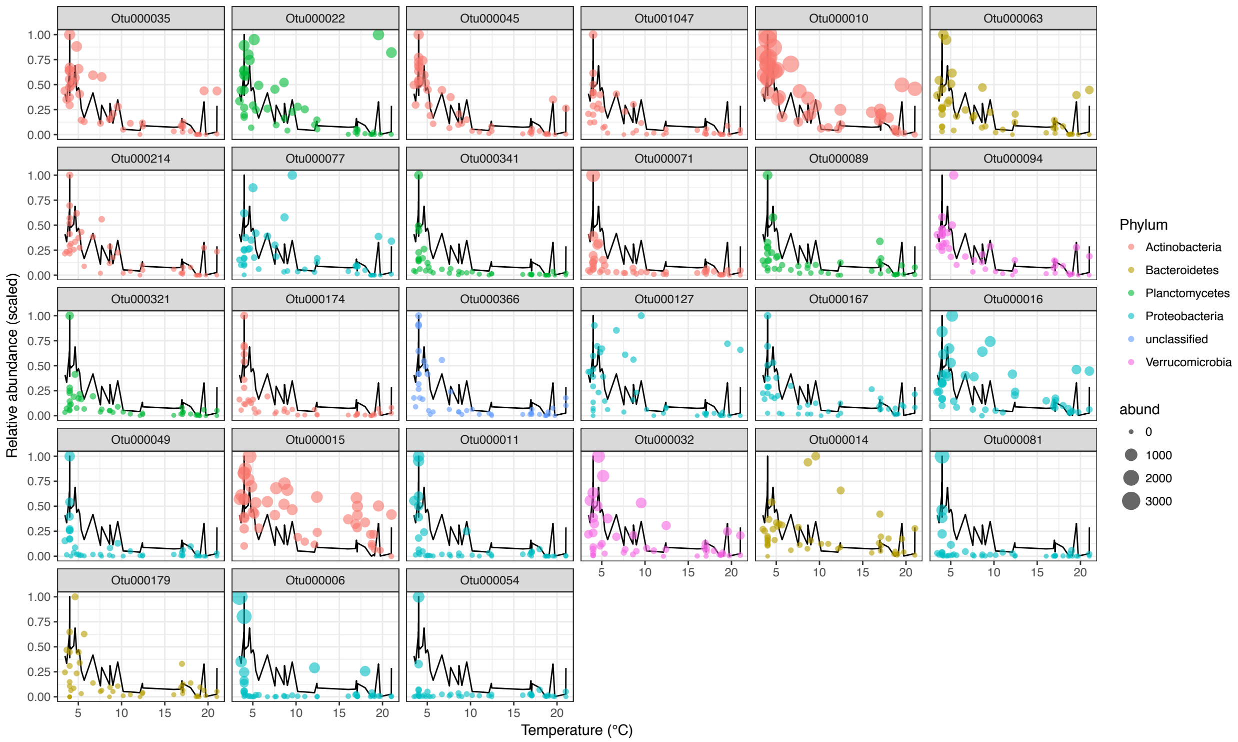


**Figure S12:** OTU abundance (scaled to 0-1 range) in the **M3** (ML module) are shown in bubbles. The PC1 trend for M3 (scaled to 0-1 range) is shown as a black line in each panel. OTU panels are arranged in order of decreasing centrality (connectedness). Bubble sizes indicate actual relative abundance values of the OTUs to communicate which OTUs were generally more or less abundant in Baikal.

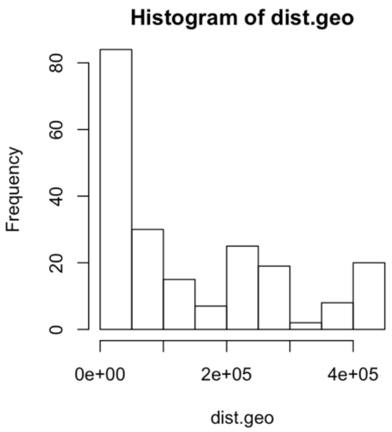


**Figure S13:** The non-normal frequency distribution of the geographic distance matrix.

**Table S1:** statistics for the four detected modules in the Baikal co-occurrence network. M1 and M3 have the highest transitivity (clustering coefficient) values.

| Module | Average Path | Transitivity |
|---|---|---|
| M1 | 1.626 | 0.684 |
| M2 | 1.797 | 0.540 |
| M3 | 1.410 | 0.823 |
| M4 | 1.795 | 0.583 |
